# From Geometry to Hierarchy: Charting the Dominant Modes of Human Cortical Function Through Development

**DOI:** 10.64898/2026.09.11.750874

**Authors:** Alexander Holmes, Kane Pavlovich, Xing Qian, Juan Helen Zhou, Chong Yap Seng, Ai Peng Tan, Michael J. Meaney, Johan G. Eriksson, Daniel S. Margulies, James C. Pang, Alex Fornito

## Abstract

Cortical function in the human cortex is spatially patterned along a macroscale information processing hierarchy. This hierarchy tracks a sensory-association axis anchored at one end by unimodal sensory regions and at the other by transmodal association areas. Biophysical models of brain dynamics suggest that this axis should align with the dominant, resonant modes of cortical anatomy and geometry, but they do not. Instead, the dominant anatomical modes correspond to simple spatial gradients extending along rostro-caudal, medio-lateral, and dorso-ventral axes.

How does this divergence between anatomical and functional modes of the cortex arise? Here, using four independent functional magnetic resonance imaging datasets spanning infancy to young adulthood (n=1,512, aged 0-21 years), we show that this spatial divergence is attributable to the maturation of long-range cortico-cortical inter-regional functional coupling. Specifically, we find that the dominant functional mode of the human cortex during the first 3 to 4 years of life closely resembles the rostro-caudal mode of cortical geometry, then shifts to a rudimentary sensory-association-like mode between 4 and 6 years of age and ultimately matures into a prototypical sensory-association mode by age 13. In infancy and early childhood, a linear model predicting the topography of the dominant functional mode from modes of cortical geometry substantially outperforms a model relying on the adult sensory-association mode, whereas the reverse is true in later stages of development. Furthermore, targeted removal of long-range connections causes the adult functional mode to regress from a sensory-association topography to an infant-like rostro-caudal mode. Our findings indicate that the maturation of long-range cortico-cortical coupling supports the gradual emergence of hierarchical modes of cortical function from an organization initially dominated by geometric constraints.

## Introduction

Cortical structure and function show a complex organization that spans scales from the cellular to the macroscopic (1,2). Recent work has shown that this complexity can be simplified using statistical decompositions that project high-dimensional anatomical or physiological data into a low-dimensional space that is defined by a small subset of latent variables corresponding to the dominant modes of the cortex (3–6). Such approaches, regardless of the details of the specific decomposition technique used, typically reveal that the leading (i.e., most dominant) mode of the human cortex corresponds to a gradient-like pattern that is anchored at one end by unimodal regions and at the other by transmodal cortices (3,5,7–9). Referred to as the sensory-association axis (7), this particular mode aligns with classical descriptions of cortical processing hierarchies (10), explains the most variance in measures of inter-regional functional coupling (FC) based on functional magnetic resonance imaging (fMRI) (3,5), and is evident in the spatial organization of intracortical myelination (6,11), cortical thickness (12), gene expression (13,14), and estimates of thalamocortical innervation (15) (for reviews, see (4,7)).

Developmental studies indicate that the sensory-association mode is not established from birth but instead comes to dominate cortical organization as the brain matures (7,8,16). Between the ages of 6 and 12 years, the sensory-association mode accounts for less variance in FC than a unimodally anchored sensorimotor mode, which extends from primary visual cortex to somatomotor and auditory areas (17,18). In adults, the sensorimotor mode corresponds to the second leading mode, accounting for less variance in FC than the primary sensory-association mode (3). This age-dependent mode reordering suggests that there is a developmental switch from sensorimotor to hierarchical modes of processing during the transition from childhood to adolescence.

Even earlier in development, studies of neonates suggest that the sensorimotor mode is still prominent, but explains nearly equivalent variance in FC to a rostro-caudal mode that corresponds to a simple linear gradient between the anterior and posterior poles of the cortex (19). This rostro-caudal mode aligns with an evolutionarily conserved gradient in the timing of regional neurogenesis and neuron density (20,21) as well as the spatial pattern of in utero cortical expansion (22). Moreover, no clear sensory-association mode has been observed in the analyses of both cortical and thalamocortical modes of neonatal FC without prior transformation to an adult template space (19,23–26), suggesting that the emergence of a sensory-association mode in childhood and adolescence follows key developmental and cognitive milestones (7,8).

What could explain the increasing dominance of the sensory-association mode through development? The early appearance of the rostro-caudal mode in neonates provides a clue. This mode aligns almost precisely with the first non-constant eigenmode of cortical geometry (27–30). Under neural field theory (NFT), an established mathematical framework for modelling macroscale brain dynamics (27,29,31–34), the geometric eigenmodes of the cortex directly correspond to the modes of its functional dynamics (29,35). This prediction arises from a general physical principle in which the dynamics of any system in the linear regime (which adequately characterizes macroscale brain dynamics; (27,36–38) can be represented in terms of the eigenmodes of its structure. Geometric modes serve this purpose in the cortex, under the assumption that cortical dynamics are dominated by travelling waves of excitation moving through a continuous cortical medium, with distinct points along the cortical sheet following a simple connectivity kernel that declines as an approximately exponential function of their distance (27,29,39). This distance-dependent connectivity aligns with the so-called exponential distance rule (EDR) that underpins many facets of mammalian connectome organization (29,32,40–42). Accordingly, recent empirical work has shown that the geometric eigenmodes of the cortex can be used to understand a diverse range of activity patterns measured with fMRI (27).

These observations suggest that the functional dynamics of the infant brain may more closely align with the assumptions of NFT; namely, that activity is largely driven by local, EDR-like, distance-dependent connectivity. We hypothesize that the gradual development of long-range connections that do not conform to a simple EDR perturbs this initial organization to yield complex spatial modes of FC that diverge from those dictated by geometry, ultimately giving rise to a sensory-association functional axis. Indeed, such long-range connections, while rare (43), are nonetheless salient aspects of mature connectomes that show a protracted period of maturation (44–47). They are generally positioned to link transmodal hubs of the cortex (48–50) and are thought to play a central role in cortical function (51,52).

Here, we test this hypothesis using four independent fMRI datasets to track the convergence between geometric and functional modes of the cortex between ages 0 and 21. We show that early brain function is strongly constrained by geometry until ages of 3–6 years, at which point a rudimentary sensory-association mode begins to emerge and continues to mature through late childhood. Using a perturbation analysis, we confirm that maturation of long-range cortico-cortical functional coupling provides a mechanism for driving the transition from geometric to hierarchical modes of cortical functional organization. Our findings thus indicate that human cortical function undergoes a gradual transition from an organization dominated by geometric constraints to one that facilitates hierarchical processing.

## Results

### Analysis overview

Our analysis used four publicly available datasets spanning infancy to young adulthood: the UNC/UMN Baby Connectome Project (BCP, n=205, 96 males, age=0–5 years), the Growing Up in Singapore Towards healthy Outcomes (GUSTO, n=233, 104 males, age=4.5 and 6 years) Project, the Nathan Kline Institute – Rockland Sample (NKI-RS, n=347, 184 males, age=6–21 years), and the Human Connectome Project – Development (HCP-D, n=650, 301 males, age=6– 21 years). For comparison against typical adult modes, we also analyzed data from the Human Connectome Project – Young Adult dataset (HCP-YA, n=994, 452 males, age=22–37 years).

For each dataset, we first estimated resting-state FC between each pair of 5,762 vertices on a cortical surface mesh representation of each hemisphere at the individual level. We then calculated a group-averaged FC matrix at separate age bands that covered key developmental milestones and also ensured approximately equal sample sizes per band: 0–2 months, 3–5 months, 6–11 months, 12–17 months, 18–35 months, and 36–60 months in the BCP dataset; 4.5 and 6 years in the GUSTO dataset; and 6–8 years, 9–10 years, 11–12 years, 13–15 years, 16–18 years, and 19–21 years in the NKI-RS and HCP-D datasets. We also obtained a group-averaged FC matrix for the adults in the HCP-YA dataset. In line with past literature, we specifically extracted the first two functional modes of these group-averaged FC matrices in each age bin using diffusion map embedding, a popular non-linear manifold learning technique (53,54) (see

Materials and Methods for details). We also estimated the first three geometric eigenmodes of cortical surface mesh representations at these same age bands using established approaches (27,55) (see Materials and Methods for details). We focus here on geometric eigenmodes due to their simplicity and direct link to NFT, providing a broader theoretical grounding for understanding the relationship between cortical structure and function.

### Functional and Geometric Modes of the Infant Cortex

We begin our analysis by considering the functional modes of infants in the BCP cohort. Across all age windows (ages 0–60 months), the primary functional mode followed a rostro-caudal pattern (Fig. 1A) that differed from the mature hierarchical sensory-association organization typically observed in adults (Fig. 1B). This mode accounted for between 20.99% and 33.43% of the variance in FC organization across the age bands (Fig. S1). Notably, the nodal line (i.e., points with zero amplitude) separating the rostral and caudal domains of the pattern shifted posteriorly with development, such that areas within the temporal lobe were grouped alongside caudal regions until 36 months and then grouped with rostral regions after this age. As a result, the primary functional mode ultimately matures to differentiate visual from non-visual regions.

**Figure 1.**
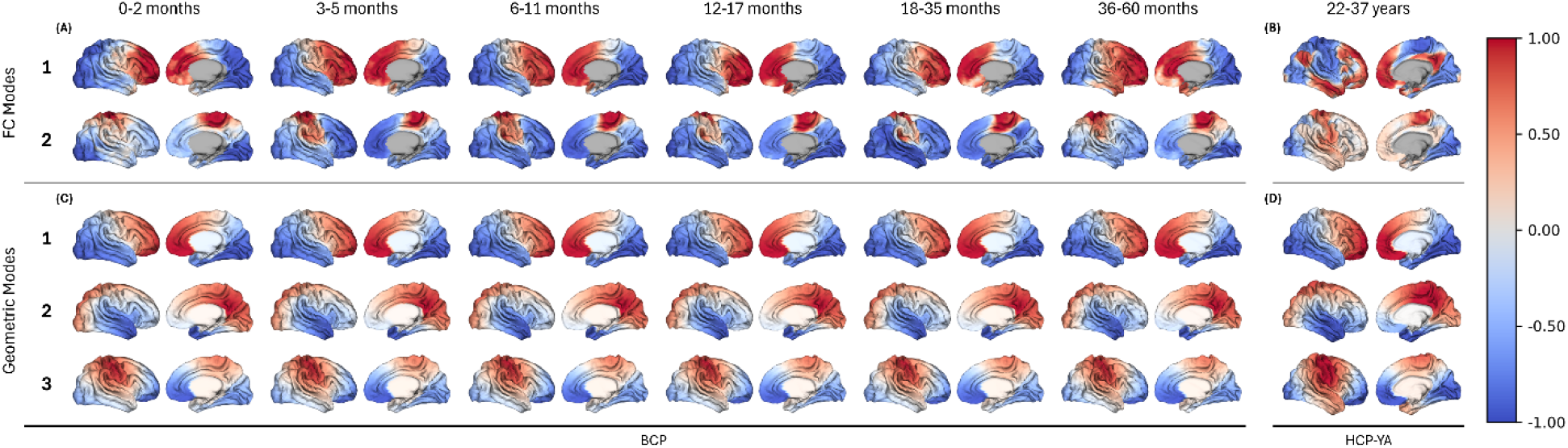
*Functional and geometric modes in infancy and adulthood.* (A) The primary and secondary functional modes (FC modes) in infancy (BCP dataset), which follow rostro-caudal and sensorimotor axes, respectively. (B) The primary and secondary functional modes in adulthood (HCP-YA dataset) instead reflect sensory-association and sensorimotor axes, respectively. (C) The first three non-constant geometric eigenmodes in infancy (BCP dataset) follow the rostro-caudal, medio-lateral, and dorso-ventral axes at all ages. (D) The first three non-constant geometric eigenmodes in adulthood (HCP-YA dataset) follow the rostro-caudal, medio-lateral, and dorso-ventral axes, similar to the infant geometric modes.

The secondary infant functional mode followed a sensorimotor pattern that corresponds to the second leading functional mode observed in adults (Fig. 1B) (3,17,19). This mode accounted for between 11.90% and 18.40% of the variance in FC organization across the age bands.

For comparison, we obtained the geometric eigenmodes of infant brains using previously described methods (27), extracted from the UNC 4D Infant Cortical Surface Atlas. The first three non-constant geometric modes followed rostro-caudal, medio-lateral, and dorso-ventral patterns (Fig. 1C), as identified previously in adults (27) (Fig. 1D). Notably, the rostro-caudal geometric mode bears a strong qualitative resemblance to the primary rostro-caudal functional mode of the infant cortex.

### Initial Signs of a Sensory-Association FC Mode in the Infant Cortex

Despite the presence of a secondary sensorimotor mode within cortical functional organization during infancy, there was no evidence of a prominent sensory-association mode within either of the first two functional modes. However, the 3^rd^ and 4^th^ functional modes both began to show initial signs of a prototypical sensory-association mode (Fig. 2A). At 0–2 months, the 3^rd^ functional mode separated temporal from non-temporal regions, while the 4^th^ mode demonstrates some minor sensory-association-like features, with a spatial correlation with the dominant sensory-association-like adult functional mode of *r* = 0.41 (Fig. 2B). However, unlike the adult sensory-association mode, the proto-sensory-association mode at 0–2 months is primarily anchored by the angular gyrus and precuneus at one end and at the other by the auditory cortex and regions of the somatomotor cortex. Although the 4^th^ mode at 0–2 months still separates other areas of the association cortex from primary sensory areas (Fig. 2A), these regions do not occupy the opposite poles of the mode, implying that they have not yet reached a sufficient degree of functional segregation. By 3–5 months, the 4^th^ mode is anchored by the visual cortex, more closely resembling the adult sensory-association mode (spatial correlation *r* = 0.48).

**Figure 2.**
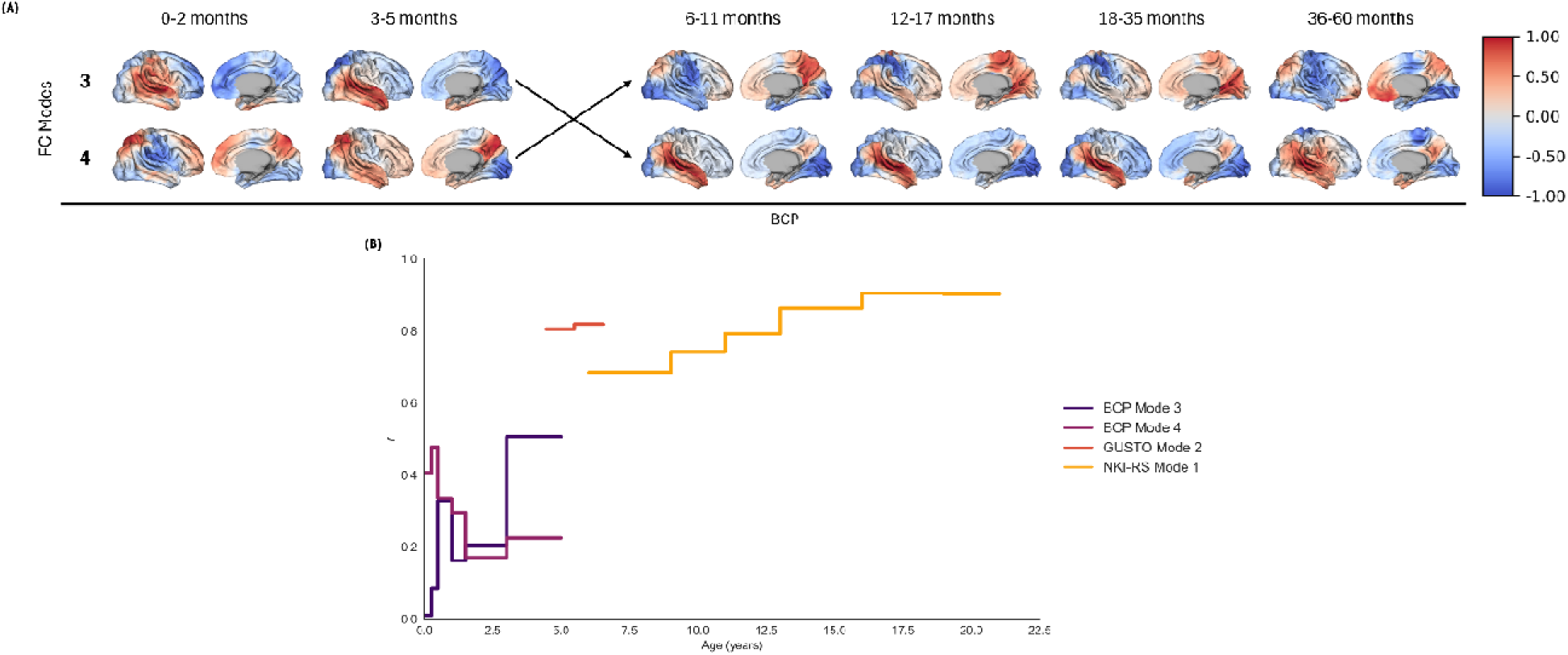
*Other lower-order functional modes reflect a proto-sensory-association mode in infancy.* (A) The third and fourth functional modes (FC modes) reflect a proto-sensory-association mode at different ages in the BCP dataset, including a developmental re-ordering between the 3–5 month and 6–11 month age bands. (B) Correlation between the adult sensory-association axis functional mode and the most “sensory-association-like” functional modes across development. The most sensory-association-like modes account for progressively increasing amounts of variance in FC with age, such that there is a switch in mode ordering from 4^th^ to 3^rd^ between 3–5 month and 6–11 month age bands, then a switch to 2^nd^ at ages 4–6 years, and then to 1^st^ from 6 years onwards.

By 6–11 months, the 3^rd^ and 4^th^ functional modes exhibit a developmental re-ordering, such that the former becomes the proto-sensory-association mode, accounting for the third largest portion of variance in FC, whereas the mode distinguishing temporal from other cortices progressively accounts for less variance. However, the proto-sensory-association mode at 6–11 months bears less of a direct resemblance to the adult sensory-association mode (0.33 ; Fig. 2B) than the 4^th^ mode at 0–2 and 3–5 months, possibly because the proto-sensory-association mode does not differentiate temporal cortex from unimodal regions. The fact that the next most-dominant mode at 6–11 months does make this differentiation implies that the elements of the proto-sensory-association axis at earlier ages split across the 3^rd^ and 4^th^ modes at 6–11 months. Accordingly, a general linear model (GLM) predicting the adult sensory-association mode using both the 3^rd^ (proto-sensory-association) and 4^th^ (temporal/non-temporal) functional modes at 6–11 months returns a similar correspondence to the sensory-association mode as previous ages(*r* = 0.48 ; Fig. S2). The re-ordering of functional modes at the 6–11 month period is likely due to a rotation of the underlying latent space, causing the 3^rd^ and 4^th^ modes to both reflect intermediate sensory-association-like states, as the structure of the low-dimensional manifold shifts across these dimensions.

In the 12–17 and 18–35 month age groups, the spatial correspondence between the 3^rd^ mode and the adult sensory-association mode decreases (*r* = 0.16 and *r* = 0.20, respectively; Fig. 2A; Fig. 2B). While the lateral surface begins to differentiate between unimodal and transmodal cortices, the medial surface remains largely undifferentiated across both age bins. Indeed, transmodal areas largely occupy a central position within the 3^rd^ functional mode, which is anchored at opposite ends by posteromedial and sensorimotor cortices. By 36–60 months, the 3^rd^ mode begins to resemble a proto-sensory-association mode that resembles the 4^th^ mode of the 0–2 and 3–5 month age bins, yielding a spatial correlation with the adult sensory-association axis of *r = 0.51* . Together, these findings suggest that transmodal areas gradually differentiate from unimodal areas between birth and around 5 years of age, but this process occurs within a latent subspace that accounts for much less of the FC variance than the rostro-caudal and sensorimotor modes.

The most sensory-association-like modes account for progressively increasing amounts of variance in FC with age, such that there is a switch in mode ordering from 4^th^ to 3^rd^ between 3–5 month and 6–11 month age bands, then a switch to 2^nd^ at ages 4–6 years, and then to 1^st^ from 6 years onwards.

### Developmental Trajectories of Functional Modes in Childhood and Adolescence

Our analysis of the BCP dataset suggests that the dominant functional mode in infants corresponds to the rostro-caudal mode of cortical geometry and that this correspondence extends until 3–5 years of age. The initial stages of a proto-sensory-association mode can be found among lower-order modes and gradually accounts for more FC variance as it begins to resemble the adult sensory-association pattern. To investigate how functional modes continue to develop beyond 5 years of age, we first investigated the GUSTO sample, which comprised two groups of children aged 4.5 years (range 4.45–4.79 years) and 6 years (range 5.48–6.55 years). As in adults (Fig. 1B), the first two modes obtained in both the 4.5 and 6 year-old age bands corresponded to the sensory-association and sensorimotor axes, but their ordering was reversed, such that the sensorimotor mode accounted for the most variance in FC organization (28.16% for 4.5 years and 32.12% for 6 years) and the sensory-association mode accounted for the second largest variance fraction (11.79% for 4.5 years and 11.11% for 6 years) (Fig. 3A) (17). This altered ordering of modes relative to adult data suggests that hierarchical cortical organization emerges at 4.5-years-old but is secondary to a dominant sensorimotor mode of organization.

**Figure 3.**
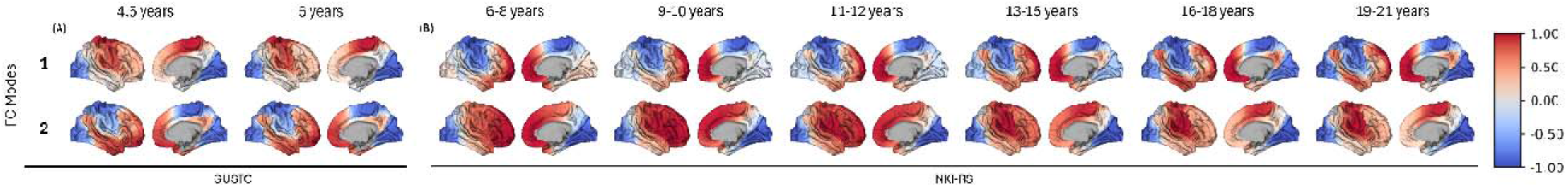
*Functional modes in early childhood and adolescence.* The primary and secondary functional modes (FC modes) in the (A) GUSTO and (B) NKI-RS datasets.

We then examined later childhood and adolescence using the NKI-RS dataset, considering participants aged 6 to 21 years. We divided the sample into 6 bins (i.e., ages 6–8, 9–10, 11–12, 13–15, 16–18, and 19–21 years) and estimated the primary and secondary functional modes separately in each bin. Within the earliest 6- to 8-year-old age band, the primary functional mode qualitatively resembled an admixture of the rostro-caudal, sensory-association, and sensorimotor patterns (Fig. 3B). The opposing poles of this mode were anchored in the frontal and somatomotor cortices, while the primary visual cortex occupied an intermediate rank. This trend continued through ages 9–10 and 11–12 years, until the dominant functional mode resembled an adult-like sensory-association mode at ages 13–15 years, with anchors localized to the visual cortex and default mode network. This change was characterized by a shift in the position of the visual network, which gradually moved closer to the somatomotor pole over time. Finally, at ages 16–18 and 19–21 years, the dominant functional mode closely resembled the sensory-association pattern that characterizes the adult functional mode, with spatial correlations exceeding .

The secondary functional mode in the 6–8-year-olds resembled the rostro-caudal mode, with a posterior midline that distinguished visual from other areas. This posterior location exhibited traits akin to the late-stage rostro-caudal mode observed in the BCP data (Fig. 1A), where the nodal line of the rostro-caudal functional mode was located at increasingly caudal positions in older age bins. After 6–8 years, regions of the somatomotor network became increasingly differentiated from other areas and gradually assumed the opposing pole to the visual cortex by 13–15 years, thus resembling the sensorimotor mode of the mature adult brain. These observations suggest that the secondary functional mode gradually transitions from rostro-caudal to sensorimotor axes between mid-childhood and peri-pubertal stages.

The developmental trends observed in the NKI-RS were partially replicated in the HCP-D sample of the same age range (Fig. S3). However, the posterior regions of the default-mode network (DMN), such as the angular gyrus, were already differentiated from somatomotor areas at 6–8 years in HCP-D participants, leading to an earlier emergence of an adult-like sensory-association pattern. Nevertheless, the visual network was still separated from the somatomotor network along the unimodal pole of the functional mode at ages 6–8, 9–10, and 11–12 years in the HCP-D cohort, with the visual network positioned closer to the unimodal anchor by age 13–15 years onwards. These results suggest that a complete differentiation of sensorimotor from association regions along the sensory-association mode may not fully emerge until mid-adolescence.

### Relative Dominance of Distinct Functional Modes Through Development

Although the spatial topographies of the primary and secondary functional modes were largely stable through infancy and early childhood, the proportion of variance in FC explained by the dominant rostro-caudal mode gradually decreased with age. During infancy in the BCP cohort, the rostro-caudal mode was most dominant at 3–5 months, explaining 33.43% of the variance in FC, before gradually receding to 20.99% by 36–60 months. Conversely, the variance explained by the sensorimotor mode increased over this same period, from 11.90% at 3–5 months to nearly equivalent to the rostro-caudal mode by 36–60 months at 18.40% (Fig. S1). These findings suggest that the first 5 years of life are associated with a declining role for the geometric rostro-caudal mode and an increasing role for the secondary sensorimotor mode in explaining the functional organization of the cortex.

At ages 4.5 and 6 years old, the dominant functional mode in the GUSTO dataset reflected a unimodal sensorimotor mode that explained 28.16% and 32.12% of the variance in FC, respectively. After 6 years of age in the NKI-RS dataset, the primary functional mode transitioned towards the sensory-association mode, which gradually accounted for an increasing fraction of variance in FC as a function of age, reaching a maximum of 37.73% at 16–18 years. The secondary sensorimotor mode explained proportionally less variance, ranging between 10% and 15% (Fig. S1). Similar trends were also observed in the HCP-D dataset, which mirrored the NKI-RS, displaying a dominant sensory-association mode and secondary sensorimotor mode at all ages (Fig. S3). These findings indicate that the sensory-association mode increasingly dominates the functional organization of the cortex from age 6 years onwards.

### Developmental Transition from Geometric to Hierarchical Modes of Brain Function

Together, the strong correspondence between FC and geometric modes from birth to 3 years of age suggests that functional brain dynamics in this developmental window are dominated by geometric constraints. However, between ages 3 and 9 years, the topography of the first two dominant functional modes is somewhat unstable, visually appearing to blend elements of the rostro-caudal, sensory-association, and sensorimotor modes before stabilizing into adult-like sensory-association and sensorimotor modes at around age 13 years. These findings suggest that the hierarchical sensory-association mode, which dominates the functional organization of the adult cortex, gradually emerges from an organization dictated by cortical geometry early in development.

To more directly quantify the emergence of hierarchical organization from geometric constraints through development, we evaluated the degree to which the first functional mode in each age bin could be reconstructed from either the first three non-constant geometric eigenmodes (i.e., the rostro-caudal, dorso-ventral, and medio-lateral modes at each age; hereafter referred to as the geometric model) or from the adult sensory-association mode (hereafter referred to as the sensory-association model) (Fig. 4).

**Figure 4.**
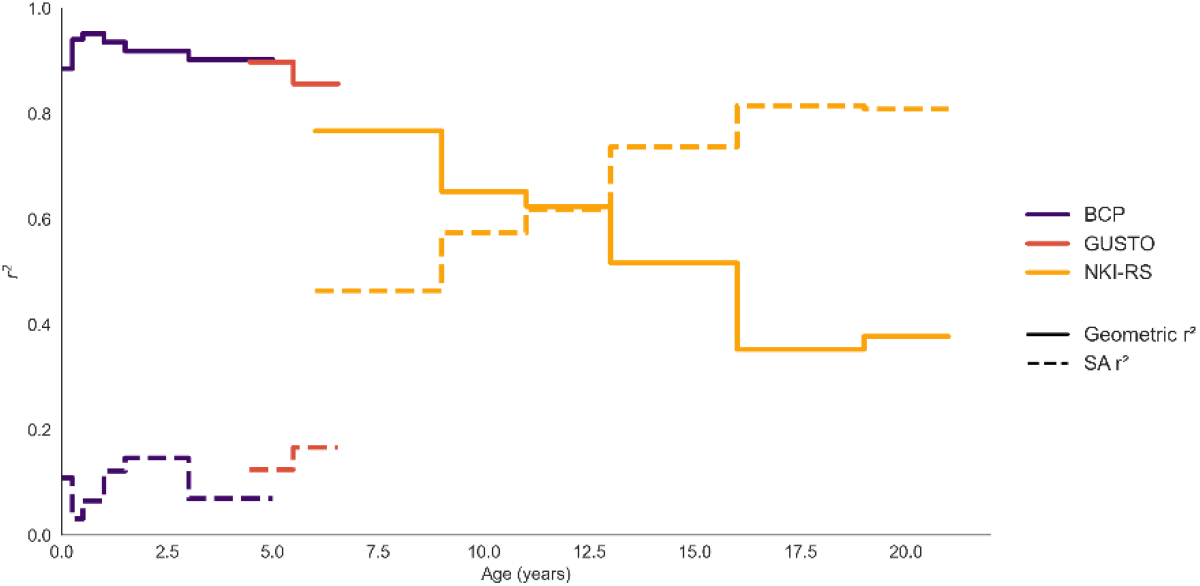
*Correspondence between age-specific functional modes and the sensory-association axis increases across lifespan.* Reconstruction of the first functional mode using either geometric or sensory-association (SA) models. The between the primary functional mode and the three geometric eigenmodes (solid lines) peaks in infancy and declines with age, while the between the primary functional mode and the sensory-association mode (dashed lines) peaks in later adolescence.

In the BCP sample, the reconstruction accuracy of the geometric model was consistently high for the primary functional mode, peaking at = 0.952 in the 6–11-month-olds and gradually decreasing with age (Fig. 4). Analysis of model parameter estimates revealed that this convergence between the functional and geometric modes in infancy was driven by the rostro-caudal spatial pattern present in the primary functional mode, which closely resembled the first non-constant geometric mode (Fig. 1; Fig. 4). In comparison, the fit of the sensory-association model in infants was poor and never exceeded = 0.146.

This trend continued in the 4.5- and 6-year-old participants of the GUSTO sample, despite the emergence of a unimodal sensorimotor mode as the dominant functional mode. The sensorimotor pattern is spatially simpler and varies over a longer wavelength than the sensory-association mode, which may explain why the geometric model, which uses only the lowest frequency geometric eigenmodes, performed well in this age group. Indeed, aspects of the sensorimotor functional mode bear a qualitative resemblance to the 3^rd^ dorso-ventral geometric mode, as both modes peak in amplitude in the somatomotor cortex. However, the dorso-ventral mode does not differentiate between the anterior prefrontal cortex and posterior visual cortex like the sensorimotor mode, highlighting how the combination of multiple geometric modes at this age contributes to the sensorimotor mode.

In the NKI-RS sample, the reconstruction accuracy of the geometric model was higher than the sensory-association model in 6–8-year-olds (*r* = 0.767 and *r* = 0.464, respectively), but the performance of the two models gradually converged and crossed over at 13–15 years, beyond which the sensory-association model showed increasingly better performance until it reached *r* = 0.808 by ages 19–21 years (compared to *r* = 0.377 for the geometric model). The increasing gap between the geometric and sensory-association models with age was replicated in the HCP-D sample (Fig. S4), but the sensory-association mode also showed superior performance at the earliest 6–8-year age band, consistent with the earlier maturation of this mode in these participants.

### Long-Range Coupling Supports the Transition from Geometric to Hierarchical Modes of Cortical Function

The strong correspondence between geometric and functional modes in infancy is consistent with the predictions of NFT and suggests that brain activity at this developmental stage is dominated by wave dynamics and EDR-like inter-regional coupling (27,29,32). To more directly test the hypothesis that the increasing divergence between geometric and hierarchical modes of function is driven by long-range cortico-cortical coupling, we divided FC estimates, measured in young adults aged 21–35 years from the HCP-YA cohort (3,56), between each pair of 5,762 cortical vertices into 10 bins according to their pairwise geodesic distance. We then re-estimated the dominant functional modes of the FC matrix after progressively removing an increasing fraction of long-range connections based on the defined bins above. In essence, this procedure reverses the age-related shift from geometry to hierarchy by investigating how long-range connections in the hierarchical mode can be perturbed to resemble the geometric mode.

We found that gradually removing an increasing fraction of longer-range connections prompted a shift in the spatial pattern of the primary functional mode from one that resembled the sensory-association mode towards the rostro-caudal mode (Fig. 5A). Specifically, sensory-association features were robust to the removal of the top 30% of the longest connections but began to resemble the rostro-caudal mode once >60% of the longest connections were removed. Supporting these trends, the correlation between the primary functional mode and the sensory-association mode decreased from = 0.994 to = 0.379 from 0% to 90% distance thresholds, respectively, whereas the correlation with the rostro-caudal mode increased from = 0.258 to = 0.952 (Fig. 5B).

**Figure 5.**
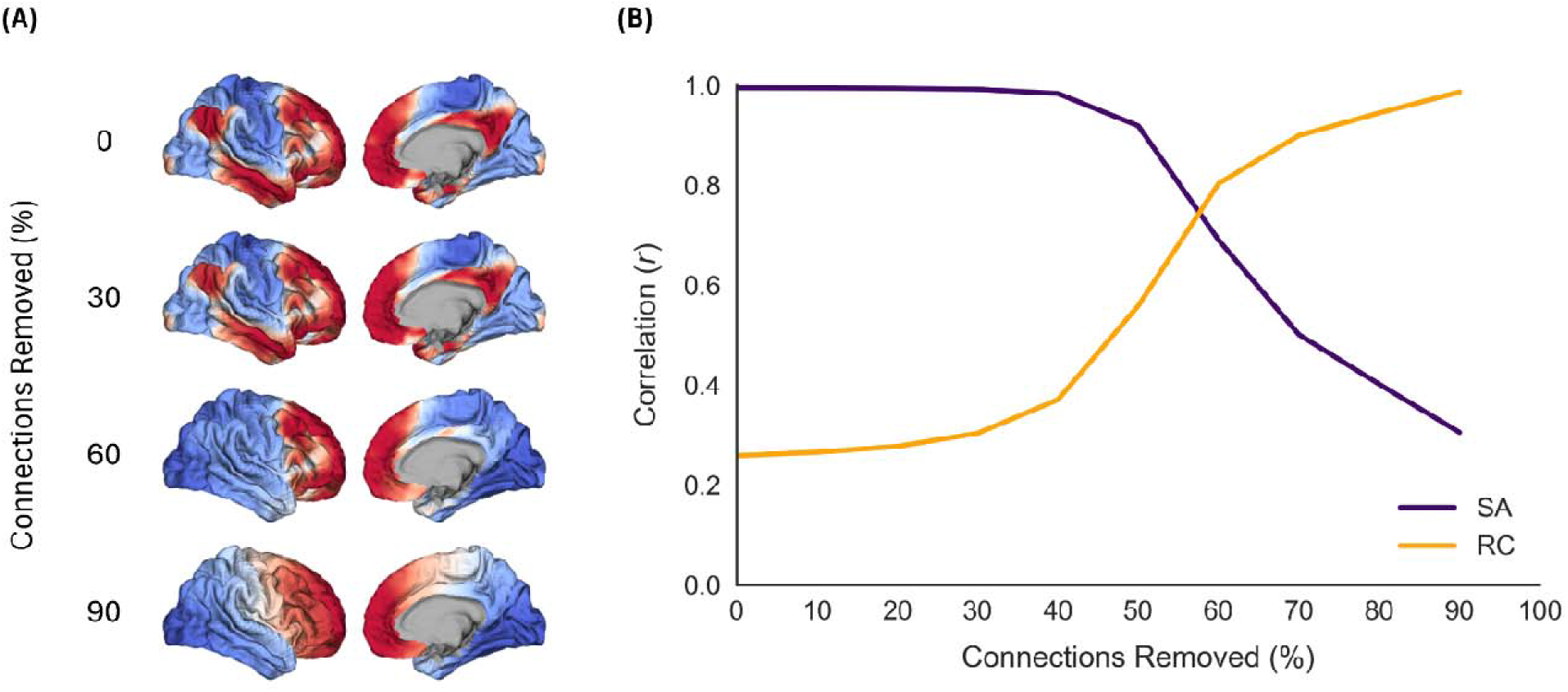
*Influence of long-distance connectivity on the dominant functional mode.* (A) The dominant functional mode obtained after removing long-range connections in the original input FC matrix, at different thresholds from 0–90% of the top distance pairs. Here we show example modes obtained at thresholds of 0%, 30%, 60%, and 90%. (B) Correlations between the dominant functional mode and either the sensory-association or rostro-caudal mode as a function of the percent of FC connections removed in decreasing order of geodesic distance. Progressive removal of an increasing fraction of long-range connections leads to the emergence of the rostro-caudal mode.

To ensure that the shift from a sensory-association to a rostro-caudal mode was driven specifically by the removal of long-range connectivity, we repeated our analyses by removing random connections with no regard to distance, with 1000 permutations of randomly selected edges. When removing edges randomly, the corresponding functional modes remained mostly unchanged, even after removing 90% of values (Fig. S5A). At this 90% threshold the average correlation between the adjusted functional mode and the sensory-association axis remained high at *r* = 0.938, while the correlation between the functional mode and the rostro-caudal mode was = 0.225 (Fig. S5B).

### Robustness Analyses

Head motion is an important confound in fMRI-based FC studies (57–59) and can vary as a function of age (60), especially in younger cohorts (61,62). To ensure that our findings cannot be explained by head motion, we repeated our BCP analyses using stricter head motion thresholds to ensure that our results are replicated in a sub-sample with low motion. Specifically, we excluded any individuals that had mean framewise displacement (FD), a summary measure of head motion (63), exceeding 0.25 mm across all time-series volumes, FD > 0.2 mm in >20% of the volumes, or FD > 5 mm in any volume, which is considered a more typical stringent threshold for motion among adult participants (57). This stricter threshold yielded a reduced sample of 147 participants. However, even within this reduced, more stringent cohort, we observed similar spatial patterns within each of the first four functional modes as seen in the present results (all), suggesting that our present results are replicated in sub-samples with low motion.

We estimated functional modes using diffusion map embedding applied to a matrix encoding inter-regional similarities in FC profiles. We used this approach to retain consistency with the bulk of the fMRI literature examining functional modes (3) but note that there are other ways of getting a low-dimensional embedding that could affect the results (64). To examine whether the specific technique used to extract functional modes affects our findings, we applied linear principal component analysis to the raw FC matrices at each age bin. In every case, and in line with prior findings (5), the first four modes obtained with linear PCA and non-linear diffusion-map embedding were highly correlated (all *r* ˃ 0.96), indicating that the choice of decomposition method does not affect our findings.

One challenge in perinatal fMRI is that participants are often asleep during scanning. This was the case for participants in the BCP, where all individuals aged under 36 months were asleep. It is thus possible that the different functional modes observed at this time represent a state-dependent effect associated with sleep, and that scanning awake infants might reveal an adult-like sensory-association mode. To evaluate the state-dependence of the spatial topography of functional modes, we performed a supplementary analysis using the ‘Simultaneous EEG and fMRI signals during sleep from humans’ dataset available from OpenNeuro (doi:10.18112/openneuro.ds003768.v1.0.10), consisting of 33 adults scanned during wakefulness and sleep (65). We found a high degree of spatial consistency in functional modes between the two states (*r* = 0.781), with the sensory-association mode being the dominant mode across all states (Fig. S6). The absence of a dominant rostro-caudal functional mode in the sleep state indicates that the infant rostro-caudal mode is unlikely to be an effect of sleep during acquisition.

## Discussion

The spatial patterning of distinct structural and functional properties along a sensory-association mode is a major organizational motif of the mature human cortex (3,4,7,66). Our analysis shows that this hierarchical organization in cortical function is not dominant from birth but instead emerges through child and adolescent development from an initial state that is largely shaped by geometric constraints. We also report evidence to indicate that the rostro-caudal mode that underpins neonatal FC is due to a relative immaturity of long-range connectivity, as eliminating these connections in adults is sufficient to trigger a transition from the sensory-association mode to a rostro-caudal architecture. Together, these findings suggest that human brain function is initially constrained by cortical geometry in early development and shifts to resemble a hierarchical sensory-association organization as long-range cortico-cortical FC matures.

### Developmental Trajectories of the Dominant Functional Modes of the Cerebral Cortex

The modes of infant FC comprised a primary rostro-caudal mode and a secondary sensorimotor mode, with no evidence for an adult-like sensory-association mode beyond a proto-sensory-association pattern distributed across the 3^rd^ and 4^th^ functional modes. The rostro-caudal mode explained proportionally less variance in FC with increasing age, before no longer being apparent in early childhood and adolescence. Previous work on neonatal functional modes has observed similar sensorimotor and rostro-caudal modes, albeit with a reversed ordering than the present study, such that the sensorimotor mode was primary and the rostro-caudal mode was secondary (19,23,67). This instability may be due to these modes accounting for similar amounts of variance in early development. Slight changes in sample composition and/or data analysis and processing may thus switch their ordering.

Although the rostro-caudal mode was consistently the most dominant functional mode in infancy, its spatial pattern shows subtle changes with age, such that the nodal line that separates the positive and negative domains of the mode shifted posteriorly with age until 36–60 months. By 36–60 months, this mode differentiated visual from non-visual regions, suggesting that the rostro-caudal mode ultimately contributes to the functional differentiation of the visual cortex along the sensorimotor axis later in development. Indeed, the infant sensorimotor mode did not show strong differentiation between visual and somatomotor cortices and explained an increasing fraction of variance in FC with age that paralleled developmental shifts in the nodal line of the rostro-caudal mode. These findings imply that differentiation along the sensorimotor axis may be a necessary precondition for the emergence of a mature sensory-association mode.

The primary mode at ages 4.5 and 6 years old within the GUSTO dataset reflected a clearer sensorimotor mode, anchored by visual and somatomotor regions at opposite poles. The geometric model nonetheless explained the dominant sensorimotor mode at these ages with high accuracy. Such high performance suggests that the sensorimotor mode may be constrained by cortical geometry in some capacity. It is thus noteworthy that the sensory areas of V1, S1/M1, and A1 are approximately positioned at the poles of the rostro-caudal, dorso-ventral, and medio-lateral geometric eigenmodes, respectively. Their spatial location may thus be dictated by geometric constraints on early cortical patterning gradients, which may prioritize the poles as target location for sensory thalamic afferents that differentiate primary unimodal from other areas (66,68,69).

The secondary functional mode in the GUSTO dataset resembled a rudimentary sensory-association axis, consistent with past work reporting a re-ordering of functional modes in childhood (17,18). Compared to the infant BCP data, children within GUSTO exhibited a higher correlation with the adult sensory-association pattern. Moreover, the functional modes representing a proto-sensory-association axis were ranked 4^th^ and 5^th^ overall during infancy, compared to 2^nd^ within early childhood. These findings suggest that the sensory-association axis progressively accounts for more variance in FC as unimodal regions functionally differentiate from transmodal regions, in line with a waning influence of geometry. Continuing this trend, the sensory-association axis dominated FC organization in later age groups, as assessed in both the NKI-RS and HCP-D datasets, accounting for an increasing fraction of FC variance, being ranked 1^st^ over the sensorimotor mode. A clear rostro-caudal pattern was not evident within the functional modes of these later age groups.

Together, our findings indicate that cortical function transitions through development, such that it is dominated by geometric constraints early on, followed by sensorimotor function, and then hierarchical processing. The early dominance of rostro-caudal and sensorimotor modes may reflect the influence of early rostro-caudal neurogenic gradients and the early maturation of sensorimotor systems (70–72). The later emergence of the sensory-association mode by around age 5 coincides with key cognitive milestones in development, such as improvements in language and basic reasoning (73,74), which are domains linked to hierarchical organization along the sensory-association axis in later life (75,76). However, it should be emphasized that developmental changes in the spatial topography of dominant functional modes are likely more complex than a simple re-ordering, being better characterized as a rotation of the low-dimensional mode space (17). Diffusion map embedding and other spectral analysis techniques identify latent manifolds within a low-dimensional space, plotting this manifold across multiple axes. Thus, for example, the re-ordering of the sensorimotor mode at around age 6 may reflect a rotation of the entire manifold across multiple axes, which may explain why the GUSTO dataset displayed a relatively “clean” primary sensorimotor mode at ages 4.5 and 6 years while the secondary sensorimotor mode in NKI-RS and HCP-D showed stronger anchoring in visual regions.

### Divergence Between Functional and Geometric Modes Increases with Age

The gradual shift from geometric to hierarchical modes of brain function that we observed could be driven by the myelination of long-range cortico-cortical or subcortical-cortical fibers in childhood (77–81). The relevance of geometric eigenmodes considered here derives from NFT, which relies on an approximation that connectivity between different cortical locations follows an isotropic, EDR-like kernel (27,29,32). In early infancy, many long-range connections are yet to fully develop, leading to a tight correspondence between geometric and functional modes, as observed here. The maturation of topologically complex connections that do not conform to this EDR may perturb the wave dynamics supported by EDR connectivity, and which are shaped by geometry (27), leading to the more complex spatial topographies evident in the dominant functional modes of the adult cortex. Our perturbation analysis aligns with this view by showing that progressive removal of long-range connections triggers a shift in the spatial architecture of the dominant functional mode in adults from the sensory-association to rostro-caudal topography. In other words, removal of long-range cortico-cortical connections is sufficient to trigger a regression from hierarchical to geometric modes of organization. These results align with evidence that many of the connections that deviate from an EDR-like trend are long-range cortico-cortical association fibers that follow a protracted period of maturation (77–80,82,83). These results are also consistent with reports that early waves of spontaneous connectivity in sensory systems are altered by thalamocortical and other afferents to yield more complex, topographically structured cortical maps (84–86).

Our findings indicate that the emergence of a sensory-association mode from geometry-driven dynamics happens sometime between 3 and 8 years, due to the absence of such patterns within the BCP cohort and the instability of the sensory-association mode in the earliest age bins of the NKI-RS and HCP-D cohorts. At age 5 years, cortical surface area and gray matter volume in prefrontal and temporal association areas increase rapidly, as connections originating from association areas mature (87,88). This period is characterized by widespread synaptic pruning (89–91) and the refinement of long-range connections between cortical areas, supporting the development of language, social cognition, and executive functions (83,92–94). Such shifts from a geometrically constrained system to a hierarchical organization may reflect the increasing capacity of transmodal association areas to efficiently integrate diverse stimuli from multiple sensory domains and anatomically disparate functional systems (75,95–99).

### Limitations and conclusions

Infant fMRI is inherently challenging because of the unique image acquisition and pre-processing required within these populations (100–102). Babies are more susceptible to head motion artefacts and can only maintain shorter scan session durations than older participants (103). Physiological attributes also bias the quality of infant neuroimaging data, as smaller brain sizes make partial volume effects more prevalent (100–102). Tissue characteristics, like water and fat content, vary dramatically with development and introduce systematic signal inhomogeneity, impacting surface reconstruction and segmentation accuracy. Although novel methodological strategies have attempted to address these effects (104), they remain difficult to fully resolve; hence, results within infant populations must be considered in relation to these issues. Similarly, infant neuroimaging often assesses sleeping populations (105,106). All infants aged 0–36 months within the BCP dataset were asleep during acquisition, while 36–60-month-old participants were given the choice to remain awake if possible. We have presented results to indicate that the spatial structure of dominant functional modes is robust across sleep and wake states in adults (Fig. S6); hence, there are no reasons to think that this consistency should differ in infancy, but the result requires confirmation in this age group.

We focused on cortico-cortical FC in this analysis, but interactions with subcortical structures are also likely to play a critical role in sculpting the modal architecture of cortical functional organization (66,69). In particular, the development of distinct corticothalamic circuits has been linked to changes in structural and functional modes of the cortex (23,67) and these circuits have developmental heterochronicities that are patterned along the sensory-association mode (8). Maturation of higher-order thalamic nuclei is likely to complement the effect of long-range corticocortical connections in supporting synchronized activity between distal cortical locations (107). Comprehensively characterizing the effects of cortical and subcortical maturation thus remains an important line of future work.

Participants older than 3 years within the BCP and younger than 8 years within the NKI-RS and HCP-D cohorts were relatively under-sampled, compared to other age bins within the respective datasets. Unfortunately, our analysis suggests that this period is perhaps the most critical in the shift from geometric to hierarchical organization of FC. Our analysis of the GUSTO sample allowed us to examine age 4.5 and 6 years and one recent study of 4-year-olds observed a dominant sensorimotor functional mode as per our analysis, suggesting that the functional constraints of geometry may decline between 3 and 5 years (108). A single dataset adequately sampling the entire age range from birth to young adulthood will be required for a fine-grained characterization of the developmental emergence of the sensory-association axis. We also emphasize that all of our datasets are cross-sectional, hence caution must be taken in interpreting the implications of our findings at the individual level.

These limitations notwithstanding, our analysis indicates that the dominant mode of cortical functional organization in humans gradually transitions from one dictated by geometric constraints to a sensory-association mode that supports hierarchical information processing, with a critical transition occurring between 4 and 8 years of age. Synthetic pruning of connections suggested that this transition is triggered by the maturation of long-range cortico-cortical connectivity. This gradual shift in dominant processing modes of the cortex may have implications for cognitive and behavioral maturation.

## Materials and Methods

### Participants

We used data from four primary resources: The UNC/UMN Baby Connectome Project (BCP), the Growing Up in Singapore Towards healthy Outcomes (GUSTO) birth cohort study, the Nathan Kline Institute – Rockland Sample (NKI-RS), and the Human Connectome Project – Development cohort (HCP-D). Table 1 outlines the key demographic information across all datasets, and Fig. S7 shows the distribution of ages in each dataset. Detailed recruitment, inclusion, and exclusion parameters for all datasets can be found in the original release papers (109–113).

**Table 1.**
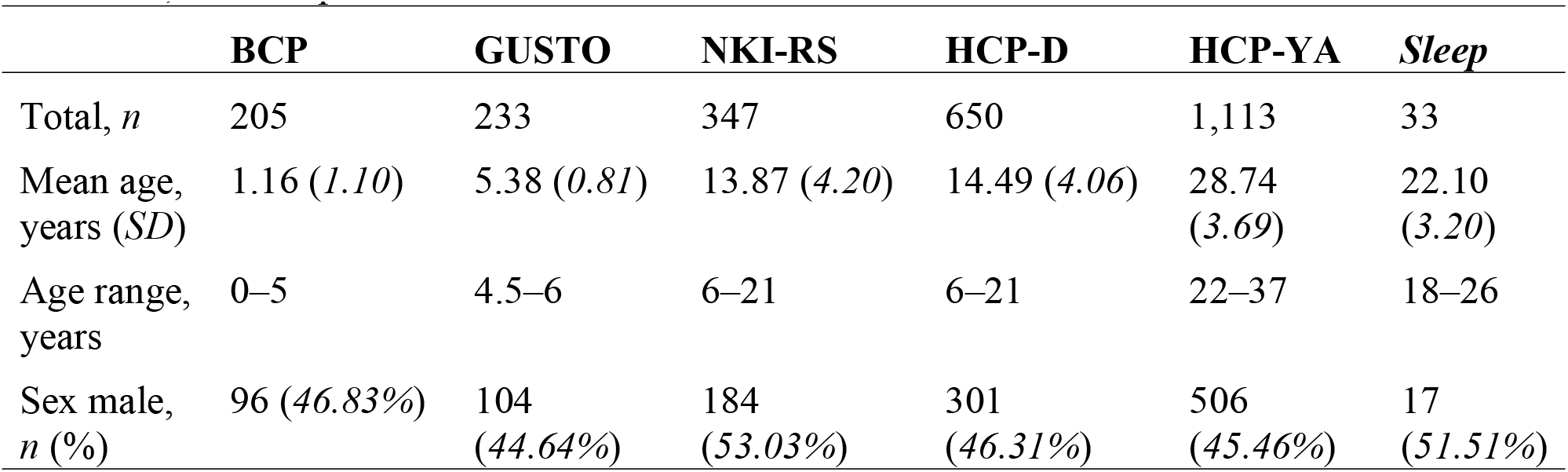
Demographic information for participants across the BCP, GUSTO, NKI-RS, HCP-D,

The BCP sample comprises infant neuroimaging data from 282 participants aged 0–60 months (110). All participants were born at a gestational age of 37–42 weeks, had a birth weight appropriate for their gestational age, and had an absence of major pregnancy and delivery complications. Following image pre-processing and various exclusions during quality control (QC) procedures, as described in the next section, we retained a final sample of 205 participants.

The initial GUSTO sample comprised 211 participants aged 4.5 years and 264 participants aged 6 years, for a total sample of 475 participants (108,112). However, following image pre-processing and QC (108), as described in the next section, we retained a final sample of 100 participants aged 4.5 years and 133 participants aged 6 years, for a total sample of 233 participants.

The full NKI-RS sample comprises 1,228 participants aged 6–85 years (111). Here, we focused only on ages 6 to 21 years due to our emphasis on investigating the emergence of an adult-like pattern of functional modes throughout childhood adolescence. Following age restrictions, image pre-processing, and QC procedures, as described in the next section, our final sample comprised 347 individuals. All participants were psychiatrically and neurologically healthy and not taking any psychotropic medication at the time of scanning.

The HCP-D sample comprises 650 participants aged 6–21 years (109,113). All participants were psychiatrically and neurologically healthy and not taking any psychotropic medication at the time of scanning. The similar age ranges of the NKI-RS and HCP-D samples allowed us to test for the reproducibility of our findings between the ages 6–21 years.

In addition to the four datasets described above, we also utilized previously published data from the Human Connectome Project – Young Adult (HCP-YA) dataset (56) to provide a baseline reference for comparisons against adult data. Here, both functional (3) and geometric modes (27) have been previously characterized in this cohort, allowing for a well-validated sample of adult cortical organization. The HCP-YA sample comprises 1,113 participants, with all participants psychiatrically and neurologically healthy and not taking any psychotropic medication at the time of scanning.

Many of the infants in the BCP cohort were sleeping at the time of scanning. To examine whether sleep could lead to the emergence of a rostro-caudal pattern as the dominant functional mode, we additionally examined a publicly available dataset of sleeping adults available from the study ‘Simultaneous EEG and fMRI signals during sleep from humans’ (65) on OpenNeuro (doi:10.18112/openneuro.ds003768.v1.0.10). This dataset comprises 33 adults, scanned during both wakefulness and sleep.

### MRI Data Acquisition and Image Pre-processing

#### UNC/UMN Baby Connectome Project (BCP) dataset

Imaging data for the BCP dataset were acquired on 3T Siemens Prisma MRI scanners at the Center for Magnetic Resonance Research at the University of Minnesota and the Biomedical Research Imaging Center at the University of North Carolina at Chapel Hill. Anatomical T1-weighted (T1w) (0.8 mm isotropic voxels, TR = 2400 ms, TE = 2.24 ms) and T2-weighted (T2w) MRI (0.8 mm isotropic voxels, TR = 3200 ms, TE = 564 ms) were acquired, in addition to at least one session of single-shot resting-state echo-planar images (EPI) MRI with AP and PA phase encoding directions (420 volumes, 5:47 minute scan length, 2.0 mm isotropic voxels, TR = 800 ms, TE = 37 ms).

Raw data were downloaded from the NIMH Data Archive and pre-processed on the MASSIVE high-performance computing facility (114) via the recently developed Nibabies pre-processing package (115). In short, anatomical images were corrected for intensity non-uniformity with N4BiasFieldCorrection (116), then skull-stripped and nonlinearly registered to the UNC 0-1-2 Infant Atlases (117) as the target template. Subject-specific brain surfaces were reconstructed using infant_recon_all within FreeSurfer 7.1.1 (104), with individual cortical surfaces manually inspected for the presence of gross artifacts and errors within reconstruction. Participants who failed reconstruction were flagged and removed from subsequent analyses.

Automatic removal of motion artifacts using independent component analysis (ICA-AROMA) (118) was performed on the pre-processed blood oxygenation level-dependent (BOLD)-fMRI data after removal of the first 10 non-steady state volumes and spatial smoothing with an isotropic, Gaussian kernel of 6 mm full-width at half-maximum (FWHM). Head-motion parameters with respect to the BOLD reference (transformation matrices and six corresponding rotation and translation parameters) were estimated before spatiotemporal filtering using MCFLIRT. Motion during fMRI acquisition was quantified by measuring framewise displacement (FD) via the Jenkinson et al. method (63). Due to the greater prevalence of motion within infant neuroimaging, participants were excluded from subsequent analyses based on the following adjusted criteria: mean FD > 0.40 mm across all timeseries volumes or FD > 7.5 mm in any volume. Following all pre-processing and image QC, 77 participants were flagged and removed from subsequent analyses, leaving a final sample of 205 participants.

Growing Up in Singapore Towards healthy Outcomes (GUSTO) dataset

Imaging data for the GUSTO dataset were acquired on a 3T Siemens Skyra scanner at KK Women’s and Children’s hospital using a 32-channel head coil. Anatomical T1w (1.0 mm isotropic voxels, TR = 2000 ms, TE = 2.08 ms) and single-shot resting state EPI MRI (116 volumes, 5:27 minute scan length, 3.0 mm isotropic voxels, TR = 2660 ms, TE = 27 ms) were acquired. Pre-processing was conducted in a previous study (108), following methods from a previous work (119). Briefly, these methods included removal of the first four frames of the resting state EPI, slice timing correction, motion correction and alignment with respect to the structural T1w image using boundary-based registration, and despiking via AFNI 3dDespike. Six motion parameters, whole brain, ventricular, and white matter signals were also nuisance regressed and removed from the data (108). Finally, participants with less than 4 minutes of scan time after removing volumes with FD > 0.6 mm or root-mean-square of voxel-wise differentiated signal >804 were excluded from analysis (108), leaving a final sample of 233 participants.

#### Nathan Kline Institute – Rockland Sample (NKI-RS) dataset

Imaging data for the NKI-RS dataset were acquired on a 3T Siemens Trio Tim scanner at Rockland County, New York. Anatomical T1w (1.0 mm isotropic voxels, TR = 1900 ms, TE = 2.52 ms) and multi-band resting-state EPI were acquired with AP and PA phase encoding directions (404 volumes, 9:35 minute scan length, 2.0 mm isotropic voxels, TR = 1400 ms, TE = 30 ms). Pre-processing was conducted in-house in accordance with the widely used HCP pre-processing pipelines (120). Basic pre-processing was conducted using FMRIB Software Library (FSL), which included skull stripping of T1w and fMRI via FSL BET, image co-registration via FSL FLIRT, and removal of the first 10 functional volumes to account for initial inhomogeneity in fMRI signals. Imaging data were nonlinearly registered to the MNI152 space with 2.0 mm resolution via FSL FNIRT. Functional data were denoised to account for motion artifacts via FSL MELODIC, which uses Independent Component Analysis (ICA) to decompose the data into distinct components corresponding to either signal or noise, whereby FSL-FIX (121,122) removed structured noise components from the functional timeseries using a hand-trained classifier from weights based on 40 manually labelled subjects randomly selected across ages 6– 21 in the NKI-RS dataset.

#### Human Connectome Project – Development (HCP-D) dataset

Imaging data for the HCP-D dataset were acquired on four 3T Siemens Prisma Scanners across Boston, Los Angeles, Minneapolis, and St. Louis. Anatomical T1w (0.8 mm isotropic voxels, TR = 2500 ms, TE = 2.22 ms), T2w images (0.8 mm isotropic voxels, TR = 3200 ms, TE = 563 ms), and at least four sessions of resting-state EPI were acquired with AP and PA phase encoding directions (488 volumes, 6:41 minute scan length for ages 8+, 3:30 minute scan length for ages 5–7, 2.0 mm isotropic voxels, TR = 800 ms, TE = 37 ms). Scan protocols and imaging software were harmonized across all sites (123,124), with one participant scanned twice at all sites to assess the influence of site effects in imaging derivatives. Preliminary analyses by the original data providers indicate that data from each site did not systematically differ in data quality metrics such as signal-to-noise, motion, or smoothness. All data were pre-processed externally by the original data providers prior to the present study, consistent with the HCP pre-processing pipelines (120). Like the NKI-RS dataset, all data were denoised through ICA and FSL-FIX (121,122) to remove structured noise components from the functional timeseries, with the accuracy of the FIX classifier visually confirmed by the original data providers.

#### Human Connectome Project – Young Adult (HCP-YA) dataset

Imaging data for the HCP-YA were acquired on a 3T Siemens Skyra Scanner at St. Louis. Anatomical T1w (0.7 mm isotropic voxels, TR = 2400 ms, TE = 2.14 ms), T2w images (0.7 mm isotropic voxels, TR = 3200 ms, TE = 565 ms), and four sessions of resting-state EPI were acquired with AP and PA phase encoding directions (1200 volumes, 14:33 minute scan length, 2.0 mm isotropic voxels, TR = 720 ms, TE = 33.1 ms). All data within the HCP-YA were pre-processed externally by the original data providers prior to the present study, following the HCP pre-processing pipelines described previously (120).

#### ‘Simultaneous EEG and fMRI signals during sleep from humans’ dataset

Imaging data for the sleeping dataset were acquired on a 3T Prisma Siemens Fit Scanner at Pennsylvania State University. Anatomical T1w (1.0 mm isotropic voxels, TR = 2300 ms, TE = 2.28ms), and two sessions of resting-state EPI were acquired with AP phase encoding direction (286 volumes, 10:00 minute scan length, 3.0 mm isotropic voxels, TR = 2100 ms, TE = 25 ms).

Multiple sessions of EPI were also obtained during sleep, with a longer scan length and the specific number of sessions varying across participants, based on their ability to remain asleep during image acquisition (286 volumes, 15:00 minute scan length, 3.0 mm isotropic voxels, TR = 2100 ms, TE = 25 ms). All data were pre-processed using fMRIprep (125) and ICA-AROMA (126).

### FC Mode Analyses

Pre-processed functional data for each participant were projected onto the individual’s cortical surface mesh, downsampled to 5,762 vertices in each hemisphere. FC matrices were then constructed by calculating pairwise product-moment correlations of vertex-wise BOLD signal timeseries. Finally, FC matrices were z-transformed and thresholded to only include the top 10% of connections at each row of the matrix, as done previously to extract the strongest pairwise connections (3,64).

In the BCP dataset, individual FC matrices were averaged across the following age bands: 0–2 months, 3–5 months, 6–11 months, 12–17 months, 18–35 months, and 36–60 months. This procedure resulted in an average FC matrix for each developmental window. Age bands were selected to ensure a minimum sample size of 20 participants within each group, while still targeting key developmental milestones. In the GUSTO dataset, FC matrices were averaged across two age bands, obtained by separating the dataset between the pre-defined 4.5-year-old and 6-year-old sample cohorts (108,112). For the NKI-RS and HCP-D datasets, FC matrices were averaged across ages 6–8 years, 9–10 years, 11–12 years, 13–15 years, 16–18 years, and 19–21 years to ensure a balanced number of participants at each developmental window.

Functional modes were estimated using diffusion map embedding (53), as implemented through the mapalign Python library (https://github.com/satra/mapalign). This approach identified a low-dimensional manifold underlying each of the age-specific FC matrices. Briefly, the method first estimates an affinity matrix of the cosine similarity between pairs of vertices in the FC matrix. Next, an eigendecomposition is applied to a diffusion probability matrix estimated from the affinity matrix, returning eigenvectors corresponding to functional modes of the FC data, ordered by the variance explained by each eigenvector.

### Geometric Eigenmode Analyses

To assess the contributions of cortical geometry to the spatial patterns of the functional modes identified through diffusion map embedding, we used the LaPy Python library (55,127) to solve the eigenvalue problem for the Laplace–Beltrami operator (LBO) of the infant, children, and adult cerebral cortices, as per past work (27). This eigenvalue problem yields the geometric eigenmodes of the cortex and is given by:

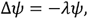

Where Δ is the LBO, which captures the intrinsic geometry of generic Riemannian manifolds such as the cortical sheet, and are the eigenmodes with corresponding eigenvalue λ. Specifically, the LBO is defined as:

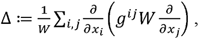

Where *x_i_*,*x_j_* are the local coordinates,*g^ij^* is the inverse of the inner product metric tensor 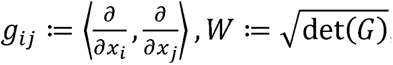, det denotes the determinant, and 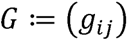.

The eigenvalue problem of the LBO is also known as the Helmholtz equation. In physically continuous systems, its solutions correspond to the spatial component of a more general wave equation and define the standing waves of the dynamics (128). Thus, under NFT, these modes correspond to the fundamental resonant modes of dynamics, or preferred patterns of excitation, much like how the musical notes played by a plucked violin string result from excitations of its eigenmodes. This is because the LBO describes the spatial propagation of the form of wave dynamics assumed by the specific version of NFT that we consider here. Detailed derivations are provided in previous work (27,29).

We estimated the LBO using a triangular surface mesh representation of the midthickness cortical surface, comprising 5,762 vertices in each hemisphere, obtained from downsampled age-specific templates from the UNC 4D Infant Cortical Surface Atlas (129). We chose this atlas to obtain a standardized representation of infant cortical architecture, benchmarked against developmental milestones. Having been developed by the original BCP data providers, and using longitudinal BCP data (110) not utilized within the present study, these template surfaces provide greater temporal continuity and are well-positioned for the present analyses. Geometric eigenmodes were obtained separately for each age band within the atlas, for ease of comparison with the similarly age-banded functional modes. Geometric eigenmodes were also obtained from a downsampled, left–right symmetric version of FreeSurfer’s fsaverage (fs_LR) population-averaged midthickness surface, also containing 5,762 vertices in each hemisphere, to be compared against the adolescent and adult functional modes. Our approach thus allowed us to investigate how changes in cortical geometry correspond with changes in functional mode architecture. Note that our analysis focused on the first three non-global geometric eigenmodes, which are largely invariant across different cortical surfaces and spherical shapes (27), and so should not depend on the specific surfaces used in this analysis.

### Characterizing Geometric Constraints and the Maturation of Functional Modes

For each age window in each dataset, we examined the degree to which the primary functional mode resembled either the geometric modes (i.e., geometric model) or the hierarchical adult sensory-association mode (i.e., sensory-association model). To assess the role of geometric constraints, we used a general linear model (GLM) that quantifies the extent to which the primary functional mode in each age bin could be explained by the first three non-global geometric modes. Note that the eigenmodes of the LBO reflect the fundamental modes of geometric variation ordered by their spatial wavelength. The first mode is always a global mode, having near constant values across the entire cortex. The next three modes describe variations along the rostro-caudal, dorso-ventral, and medio-lateral axes; progressively higher-order modes describe geometric variations occurring over higher spatial frequencies or shorter wavelengths. The lowest wavelength modes are the least physically stable and most likely to be excited (130,131). Given that many genes influencing early brain development are expressed along the rostro-caudal, dorso-ventral, and medio-lateral axes (68,69,132), we focused principally on these modes. We also quantified the similarity of the primary functional mode with the adult sensory-association mode using a separate GLM. The adult mode was obtained in an independent sample of healthy adults, corresponding to the mature sensory-association mode, as provided by the HCP-YA dataset (56). For both geometric and sensory-association models, model fit was assessed using the coefficient of determination ().

### Assessing the Role of Long-Range Connectivity

To assess the role of long-range connectivity in the emergence of a dominant sensory-association functional mode, we obtained functional modes in an adult population using the HCP-YA dataset (56). Using a matrix of pair-wise geodesic distances between vertices in the fs_LR template surface, we identified the most distant pairs of regions in each row. The FC matrix was thresholded to progressively remove these long-range connections. We started by excluding the top 10% most distant pairs and repeated the process, in increments of 10%, until the top 90% of the most distant pairs were removed. At each increment, we then retained only the 10% strongest connections for each row of the FC matrix, as done previously (3,64) and in our main analysis, to ensure that an equivalent number of datapoints were included in the embedding, regardless of any distance-based threshold. We then applied diffusion map embedding to the resulting matrices to obtain the new functional modes. This procedure thus allowed us to evaluate how the progressive removal of long-range connections changes the architecture of functional modes of the cerebral cortex. We then compared the results of this approach with an alternative perturbation strategy, which removed connections randomly with no regard to distance across 1000 permutations of randomly selected edges.

## Acknowledgments

This work was supported by Monash eResearch capabilities, including the M3 MASSIVE high-performance computing facility.

## Author contributions

Conceptualization: AH, JCP, AF

Methodology: AH, JCP, AF

Software: AH, KP, JCP

Investigation: AH

Resources: AH, XQ, JHZ, AF

Data Curation: AH, KP, XQ

Supervision: AF, JCP

Writing—original draft: AH

Writing—review & editing: AH, KP, XQ, JHZ, DM, JCP, AF

## Data and materials availability

All data are available in the main text or the supplementary materials, with the raw BCP, GUSTO, NKI-RS, and HCP-D datasets available online from the original data providers.

