## Supplementary Materials for "From Geometry to Hierarchy: Charting the Dominant Modes of Human Cortical Function Through Development"

Alexander Holmes* *et al.*

**This PDF file includes:**

Figs. S1 to S7

Fig. S1.
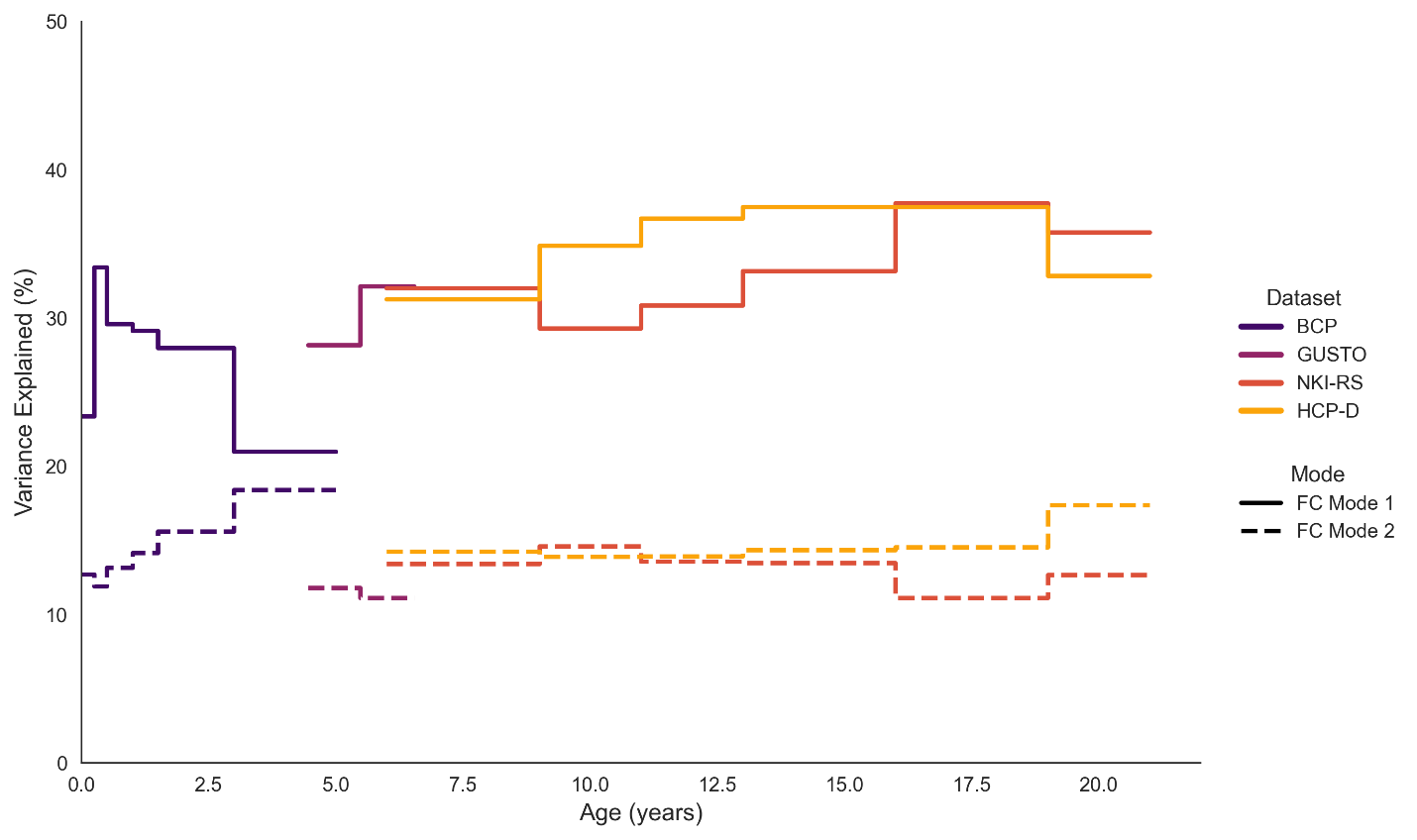

*Proportion of variance explained by the first two dominant functional modes across the lifespan.* Solid lines: Functional Mode (FC Mode) 1; dashed lines: FC Mode 2; across the BCP (indigo), GUSTO (purple), NKI-RS (red), and HCP-D (yellow) datasets.

Fig. S2.

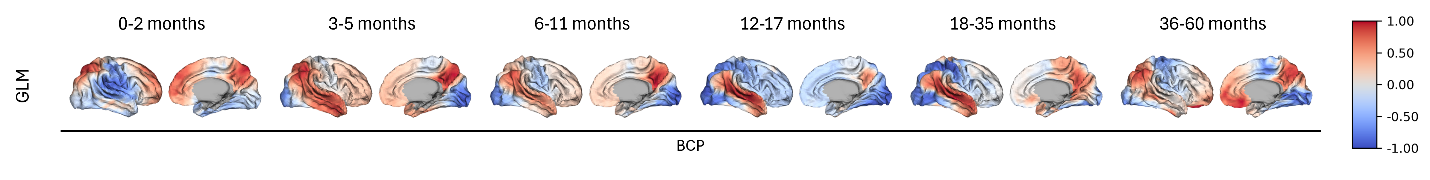

*General linear model (GLM) prediction of the adult sensory-association mode from the 3rd and 4^th^ functional modes in infancy.*

Fig. S3.

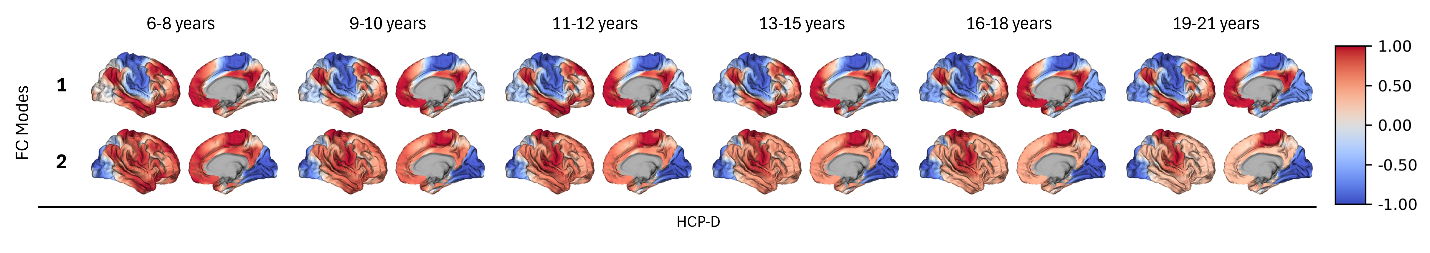

*Functional modes in early childhood and adolescence.* Spatial topography of the primary (top row) and secondary (bottom row) functional modes (FC modes) in the HCP-D dataset.

Fig. S4.

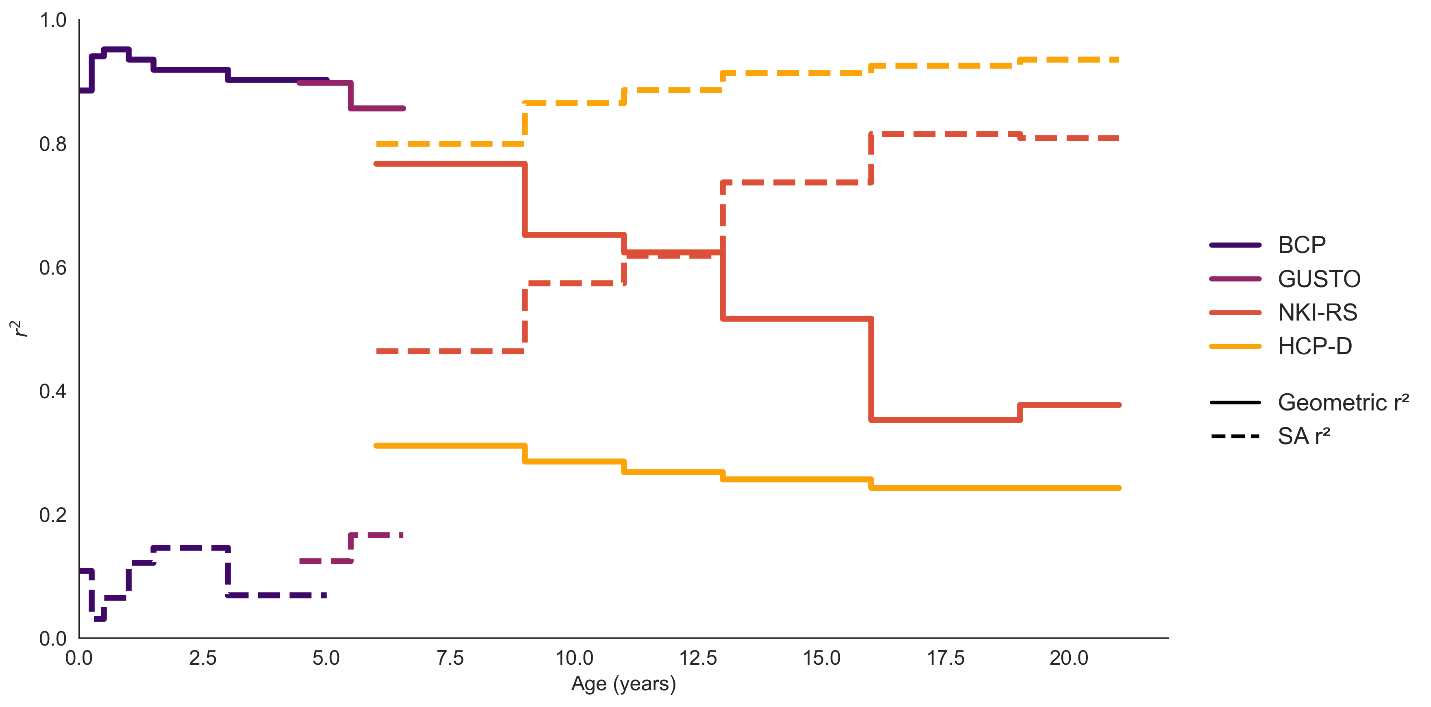

*Correspondence between age-specific functional modes and the sensory-association axis increases across the lifespan, including the additional HCP-D cohort.*

Fig. S5.

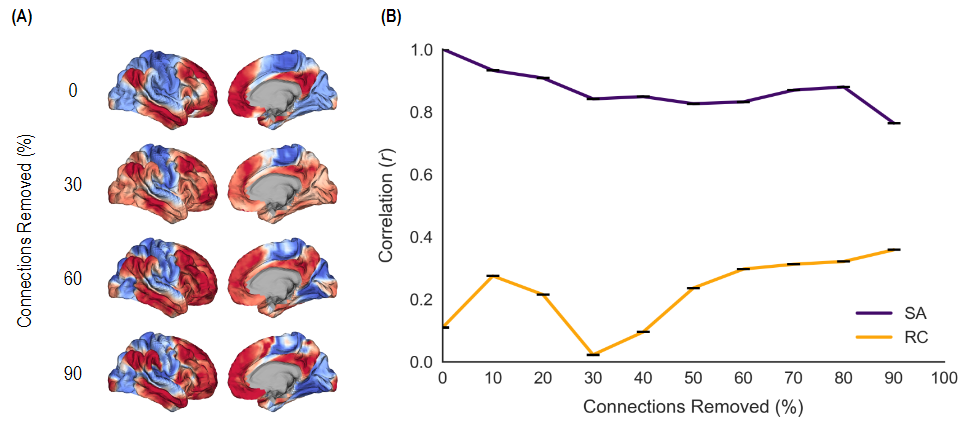

*Effect of removing random connections on the dominant functional mode.* (A) The dominant functional mode (FC mode) upon masking random connections in the original input FC matrix, at different thresholds from 0–90% of randomly chosen connections. (B) Correlation between the dominant FC mode after removing random connections with the SA and RC modes. Progressive removal of an increasing fraction of random connections does not lead to the emergence of the RC mode.

Fig. S6.

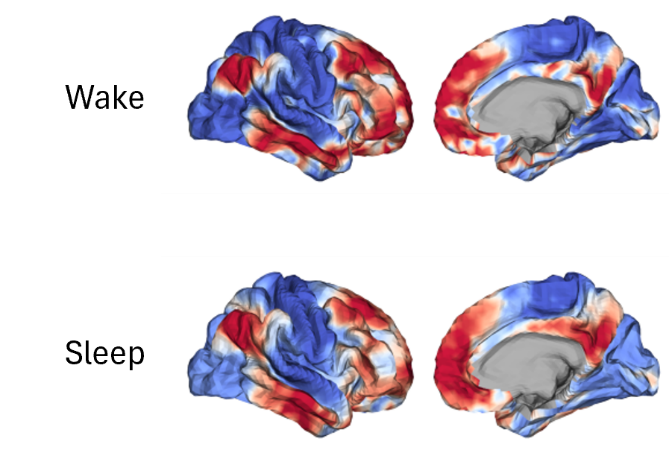

*Functional modes in wake and sleep.* The primary functional mode displays a similar spatial pattern across both wake and sleep states, reflecting a dominant sensory-association axis.

Fig. S7.

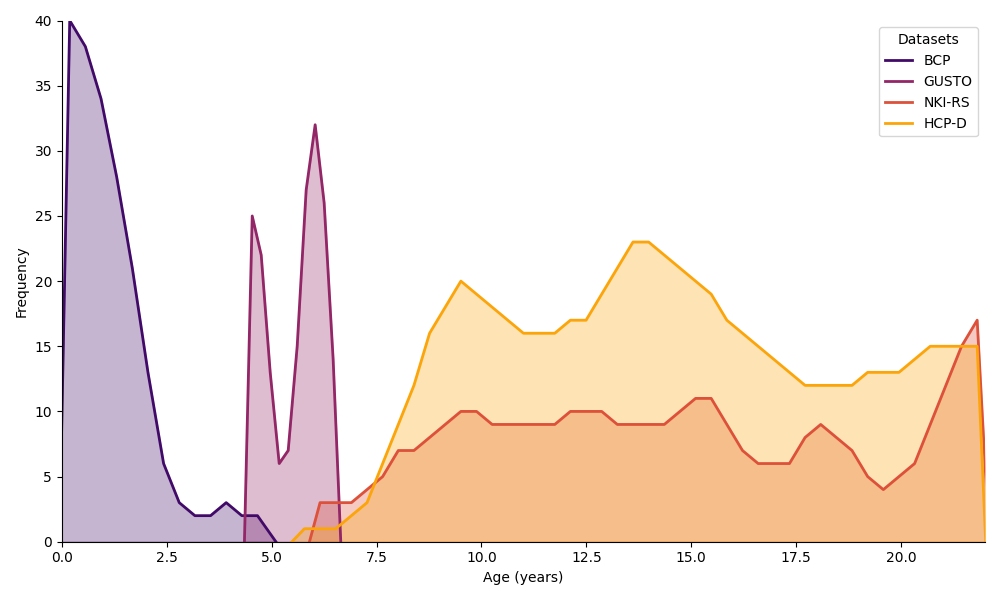

*Distribution of ages across the BCP (indigo), GUSTO (purple), NKI-RS (red), and HCP-D (yellow) datasets.*
